# The genotypic and genetic diversity of enset (*Ensete ventricosum*) landraces used in traditional medicine is similar to the diversity found in starchy landraces

**DOI:** 10.1101/2020.08.31.274852

**Authors:** Gizachew Woldesenbet Nuraga, Tileye Feyissa, Kassahun Tesfaye, Manosh Kumar Biswas, Trude Schwarzacher, James S. Borrell, Paul Wilkin, Sebsebe Demissew, Zerihun Tadele, J.S. (Pat) Heslop-Harrison

## Abstract

**Background:** Enset (*Ensete ventricosum*) is a multipurpose crop extensively cultivated in southern and southwestern Ethiopia for human food, animal feed and fiber. It contributes to the food security and rural livelihoods of 20 million people. Several distinct enset landraces are cultivated for their uses in traditional medicine. Socio-economic changes and the loss of indigenous knowledge might lead to the decline of important medicinal landraces and their associated genetic diversity. However, it is currently unknown whether medicinal landraces are genetically differentiated from other landraces. Here, we characterize the genetic diversity of medicinal enset landraces to support effective conservation and utilization of their diversity

**Results:** We evaluated the genetic diversity of 51 enset landraces of which 38 have reported medicinal value. A total of 38 alleles were detected across the 15 SSR loci. AMOVA revealed that 97.6% of the total genetic variation is among individual with an F_ST_ of 0.024 between medicinal and non-medicinal landraces. A neighbor-joining tree showed four separate clusters with no correlation to the use values of the landraces. Principal coordinate analysis also confirmed the absence of distinct clustering between the groups, showing low differentiation among landraces used in traditional medicine and those having other use values.

**Conclusion:** We found that enset landraces were clustered irrespective of their use value, showing no evidence for genetic differentiation between enset grown for ‘medicinal’ uses and non-medicinal landraces. This suggests that enset medicinal properties may be restricted to a more limited number of genotypes, a product of interaction with the environment or management practice, or partly misreported. The study provide baseline information that promotes further investigations in exploiting the medicinal value of these specific landraces

## 1. Introduction

Enset (*Ensete ventricosum;* also called Abyssinian banana) is a herbaceous, monocarpic perennial plant that grows from 4 to 10 m in height. It resembles and is closely related to bananas in the genus *Musa*, and these together with the monotypic genus *Musella* form the family *Musaceae* [1]. Enset is a regionally important crop, mainly cultivated for a starchy human food, animal feed and fiber. It contributes to the food security and rural livelihoods of 20 million people in Ethiopia. It is resilient to extreme environmental conditions, especially to drought [2] and it is considered as a priority crop in areas where the crop is grown as a staple food [3].

The use of indigenous plant species to treat a number of ailments such as cancer, toothache and stomach ache in different parts of Ethiopia has been frequently reported [4–6]. In addition to the extensive use of enset as human food and animal feed, some enset landraces play a well-known and important role in traditional medicine due to their use in repairing broken bones and fractures, assisting the removal of placental remains following birth or an abortion, and for treatment of liver disease [7–9], In the comparison of different medicinal plant species, *Ensete ventricosum* ranked first by the local people [10]. Micronutrient composition analysis of enset landraces indicates that high arginine content could be one reason for their medicinal properties, as it is associated with collagen formation, tissue repair and wound healing via proline, and it may also stimulate collagen synthesis as a precursor of nitric oxide [11].

The loss of some valuable enset genotypes due to various human and environmental factors has been previously reported [12, 13]. Medicinal landraces may be more threatened than others because when a person is ill, the medic usually gets given the plant (free of charge) to cure the ailment of the patient, but the farmer does not have an economic reason to propagate and replant the medicinal landraces. Moreover, of these landraces are highly preferred by wild animals like porcupine and wild pig [14] and more susceptible to diseases and drought [15]. Since these factors might lead to the complete loss of some of these important landraces, attention needs to be given to the conservation and proper utilization of the landraces that play important roles in traditional medicine.

The most common methods of conserving the genetic resources of vegetatively propagated plants like enset is in a field gene bank, which is very costly in terms of requirements for land, maintenance, and labor. In such cases, a clear understanding of the extent of genetic diversity is essential to reduce unnecessary duplication of germplasm [16]. Assessment of diversity using phenotypic traits is relatively straight forward and low cost [17], and is the first step in identifying duplicates of accessions from phenotypically distinguishable cultivars. However, due to the influence of environment on the phenotype, evaluating genetic variation at molecular level is important.

Molecular markers are powerful tools in the assessment of genetic diversity which can assist the management of plant genetic resources [18, 19]. Previous enset genetic diversity studies have used molecular techniques including amplified fragment length polymorphism (AFLP) [13], random amplification of polymorphic DNA (RAPD) [20], inter simple sequence repeat (ISSR) [21] and simple sequence repeat (SSR) [22]. SSR markers are highly polymorphic, co-dominant and the primer sequences are generally well conserved within and between related species [23]. Recently, Gerura *et al*. [24] and Olango *et al*. [25] have reported measurement of genetic diversity of enset using SSR markers. The previous studies were carried out on landraces from specific locations, and there was no identification and diversity study on enset landraces used for traditional medicine, potentially another reservoir of diversity. Therefore, the current study was conducted to investigate the extent of genetic diversity and relationship that exists among enset landraces used in traditional medicine.

## 2. Materials and Methods

### 2.1 Plant material and genomic DNA extraction

Thirty eight cultivated and named *Ensete ventricosum* landraces used in the treatment of seven different diseases were identified with the help of knowledgeable village elders from 4 locations (administrative zones) of north eastern parts of SNNP regional state of Ethiopia (Figure 1). For comparison, 13 enset landraces that have other non-medicinal use values were also included (principally used for human food), which brought the total number of landraces to 51. The samples were collected from individual farmers’ fields, located at 18 Kebele (the lowest tier of civil administration unit) from across the enset distribution. Since different landraces may have been given the same vernacular name at different locations [25], landraces having identical names, but originated from different locations, were labeled by including the first letter of names of location after a vernacular name. To test the consistency of naming of landraces within each location, up to 4 duplicate samples (based on their availability) were collected for some landraces, so that a total of 92 plant samples were collected (Table 1). Healthy young cigar leaf (a recently emerged leaf still rolled as a cylinder) tissue samples were collected from individual plants from November to March 2017 and they were stored in a zip locked plastic bags containing silica gel and preserved until extraction of genomic DNA. The dried leaf samples were ground and genomic DNA was isolated from 150 mg of each pulverized leaf sample following modified CTAB (cetyltrimethylammonium bromide) extraction protocol [26].

**Table 1.**
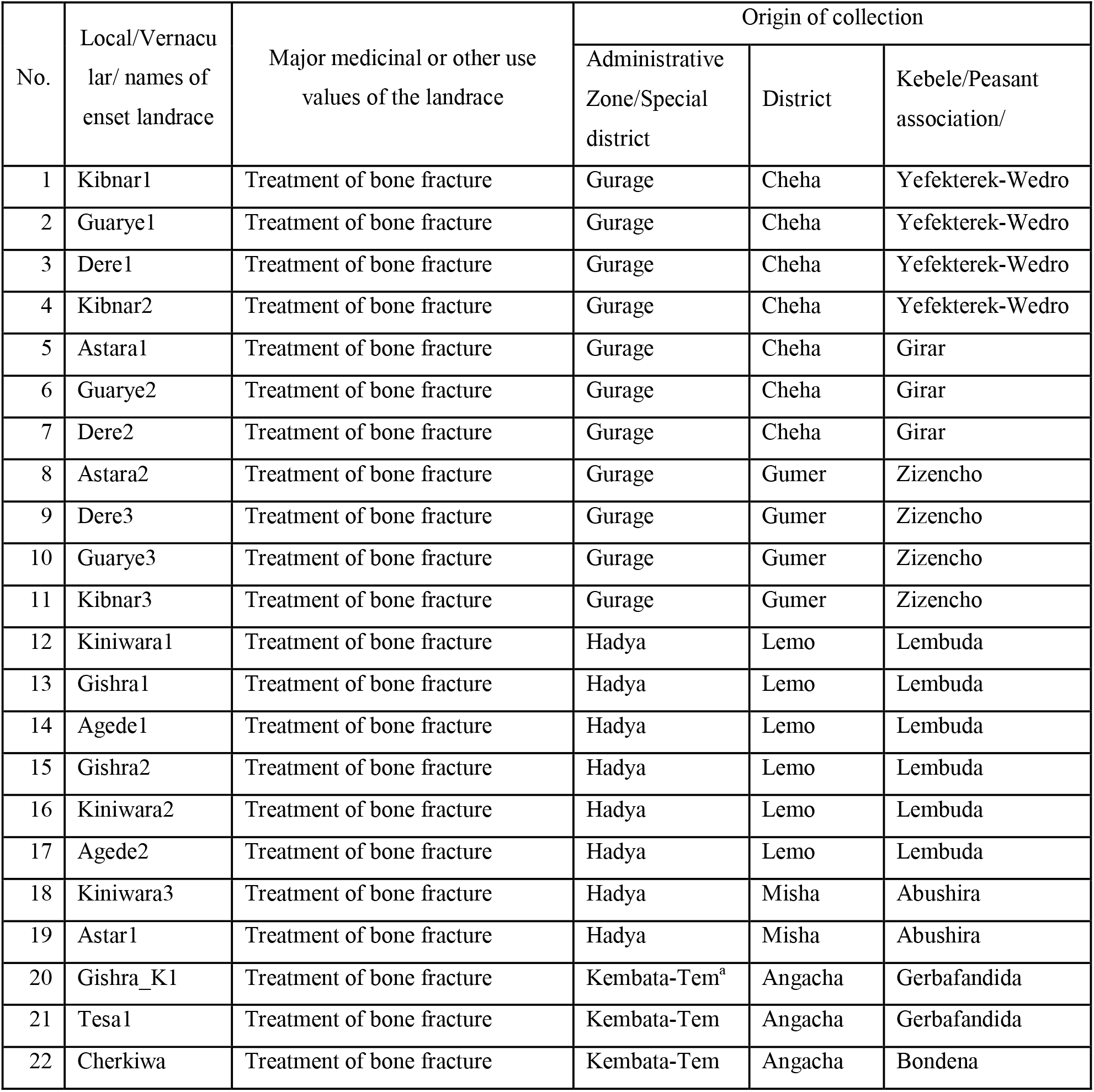
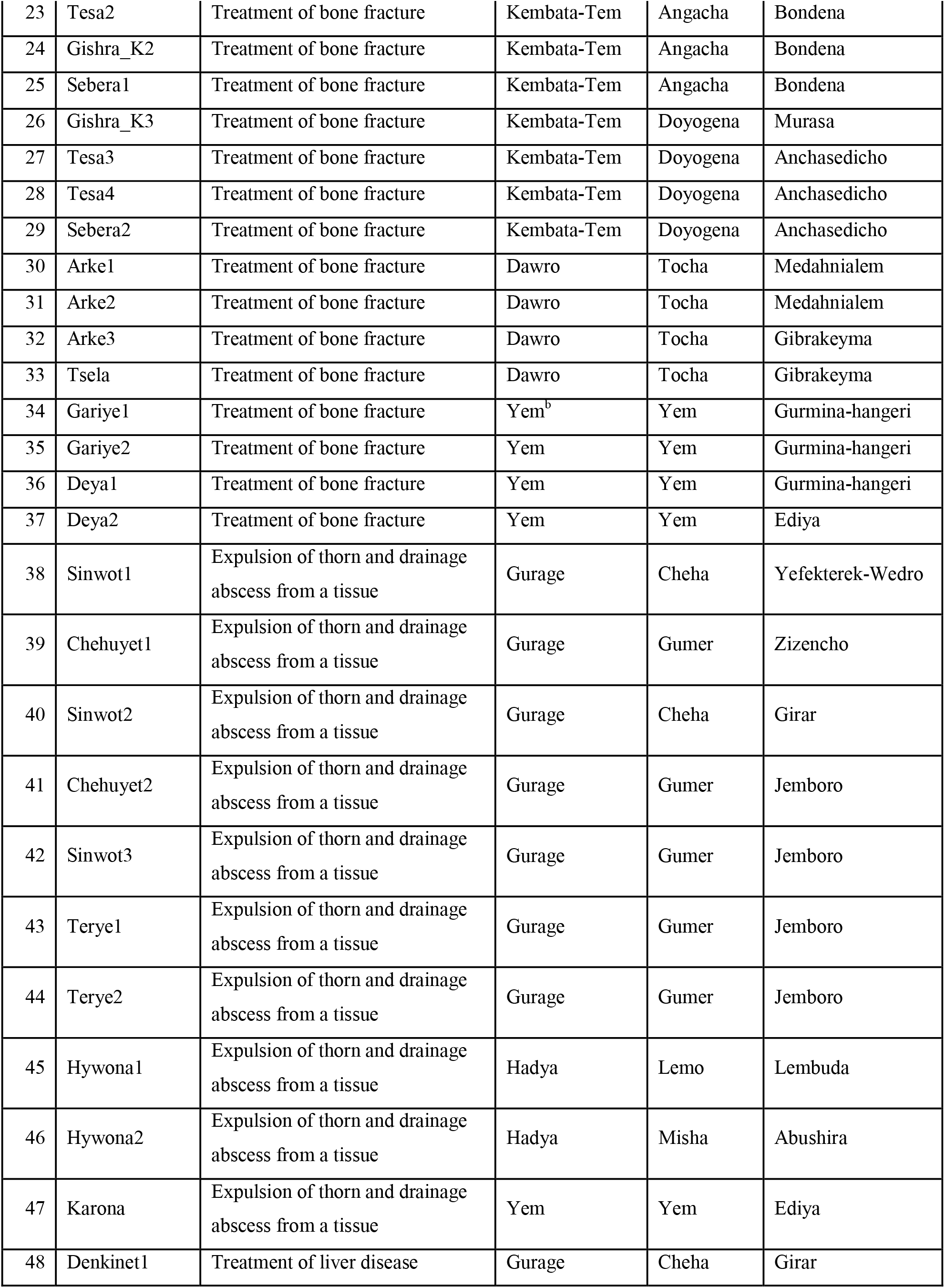
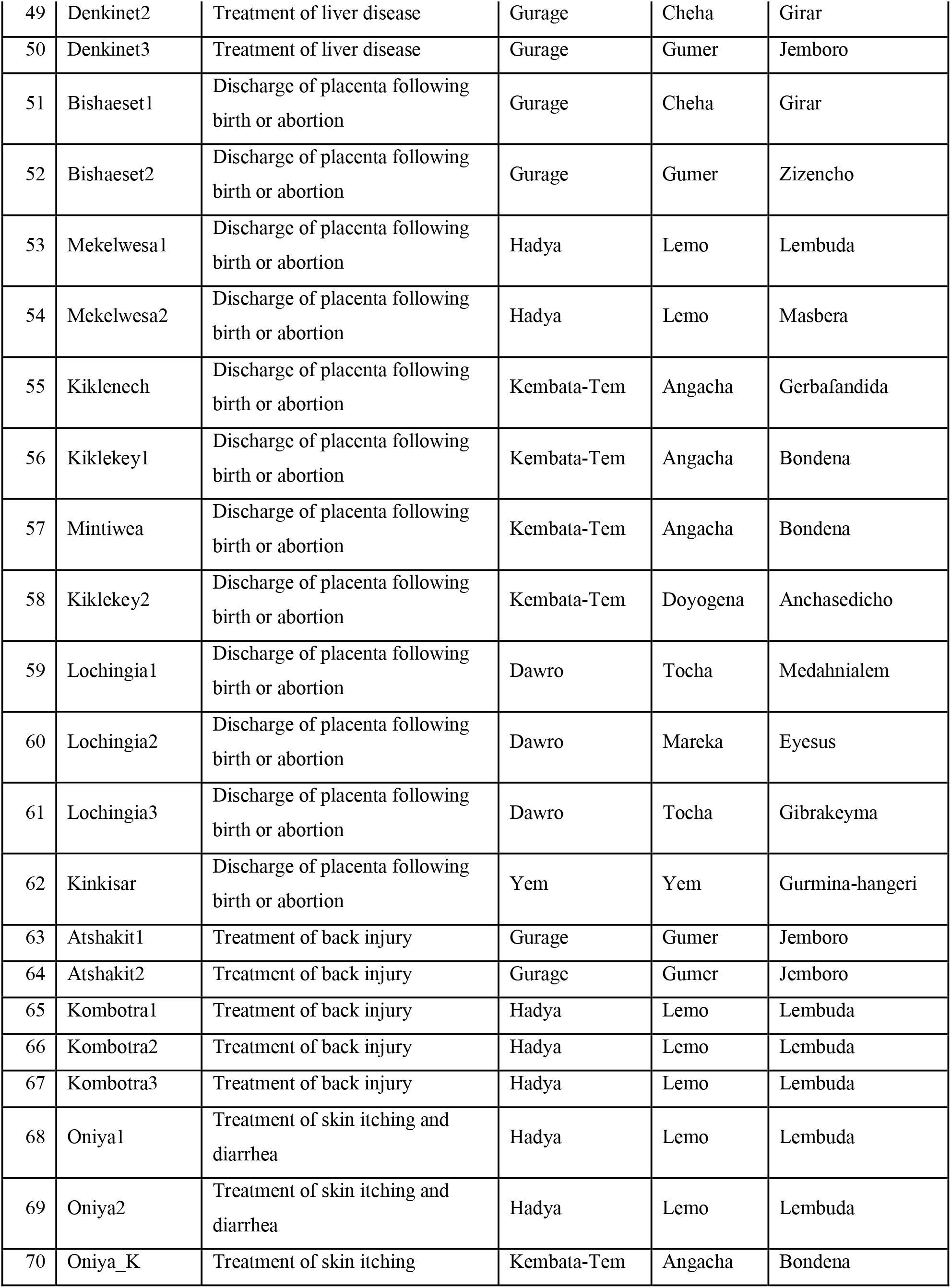
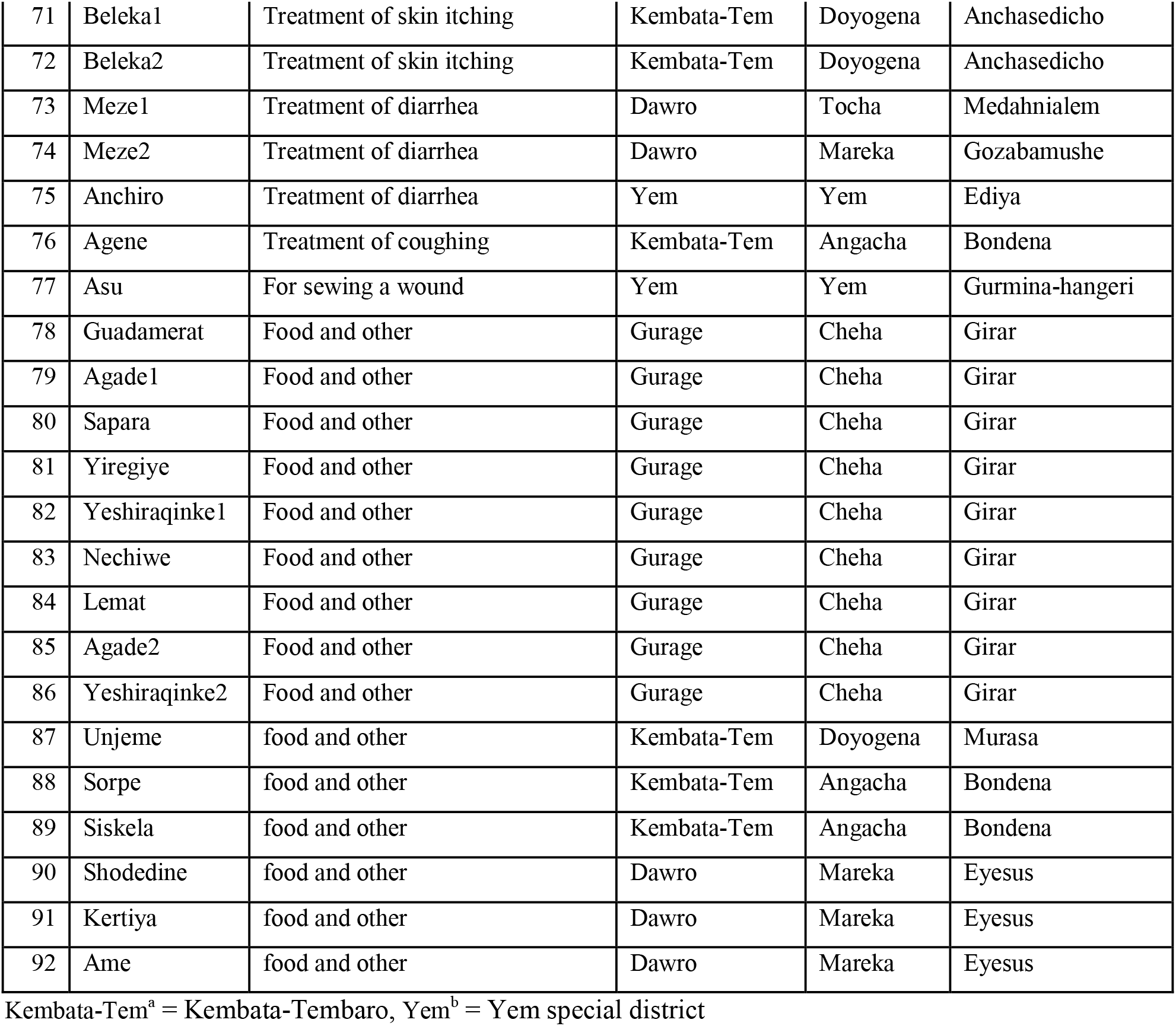
Description of plant materials used in genetic diversity study of enset (*Ensete ventricosum*) landraces using SSR markers

**Figure 1.**
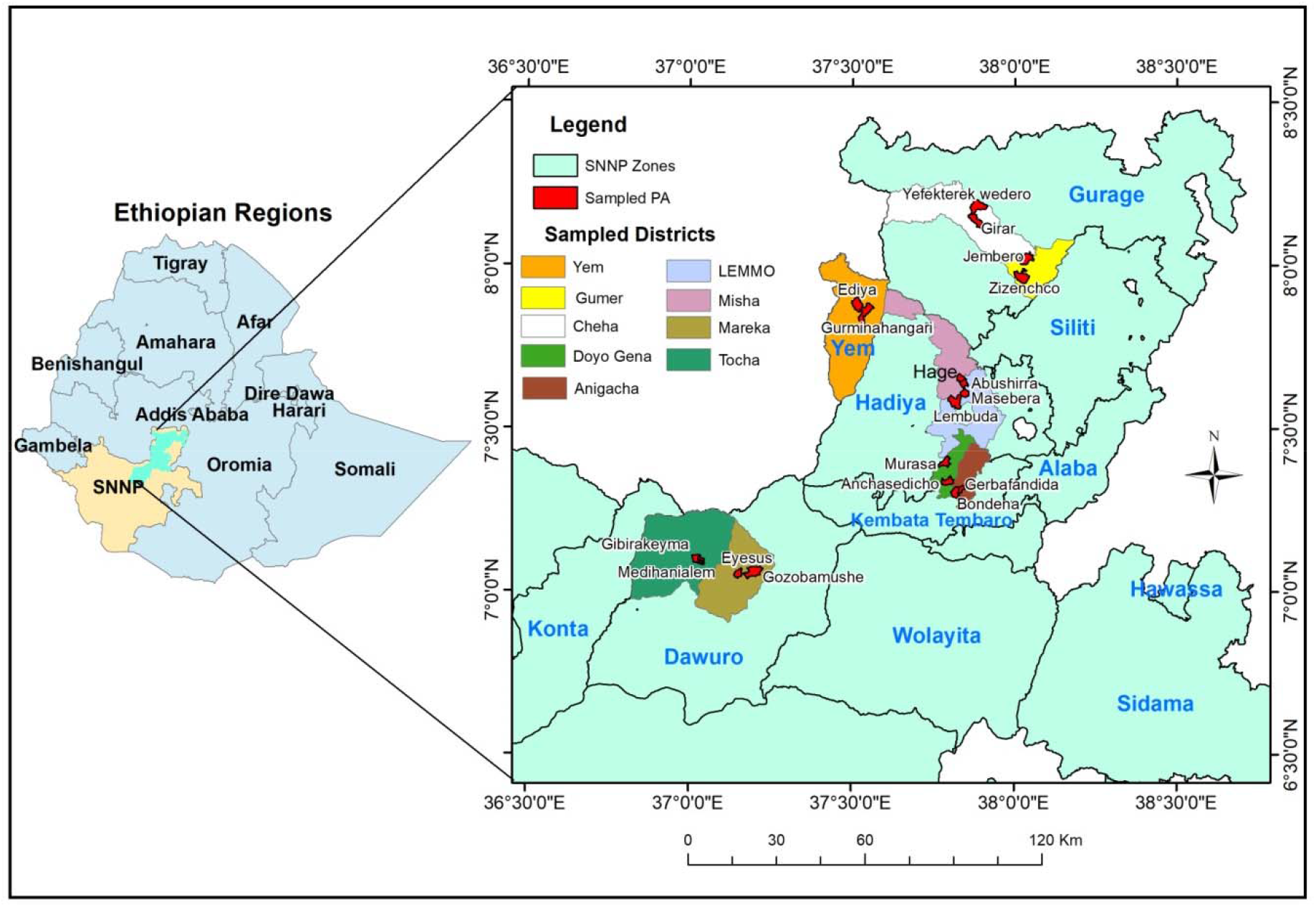
Map of Ethiopia showing its Federal Regions (left) and enset sample collection sites that represent the 9 studied districts found in, within 4 zones and 1 special district of SNNP Region. The map was constructed using geographic coordinates and elevation data collected from each sites using global positioning system (GPS). PA= Peasant association (Kebele)

### 2.2 PCR amplification and electrophoresis

Twenty one enset SSRs primer-pairs (14 from Olango *et al*. [25], and 7 from Biswas *et al*., (unpublished and http://enset-project.org/EnMom@base.html)) were initially screened for good amplification, polymorphism and specificity to their target loci using 15 samples. This led to the selection of 15 primer-pairs to genotype the landraces (Table 2).

**Table 2.**
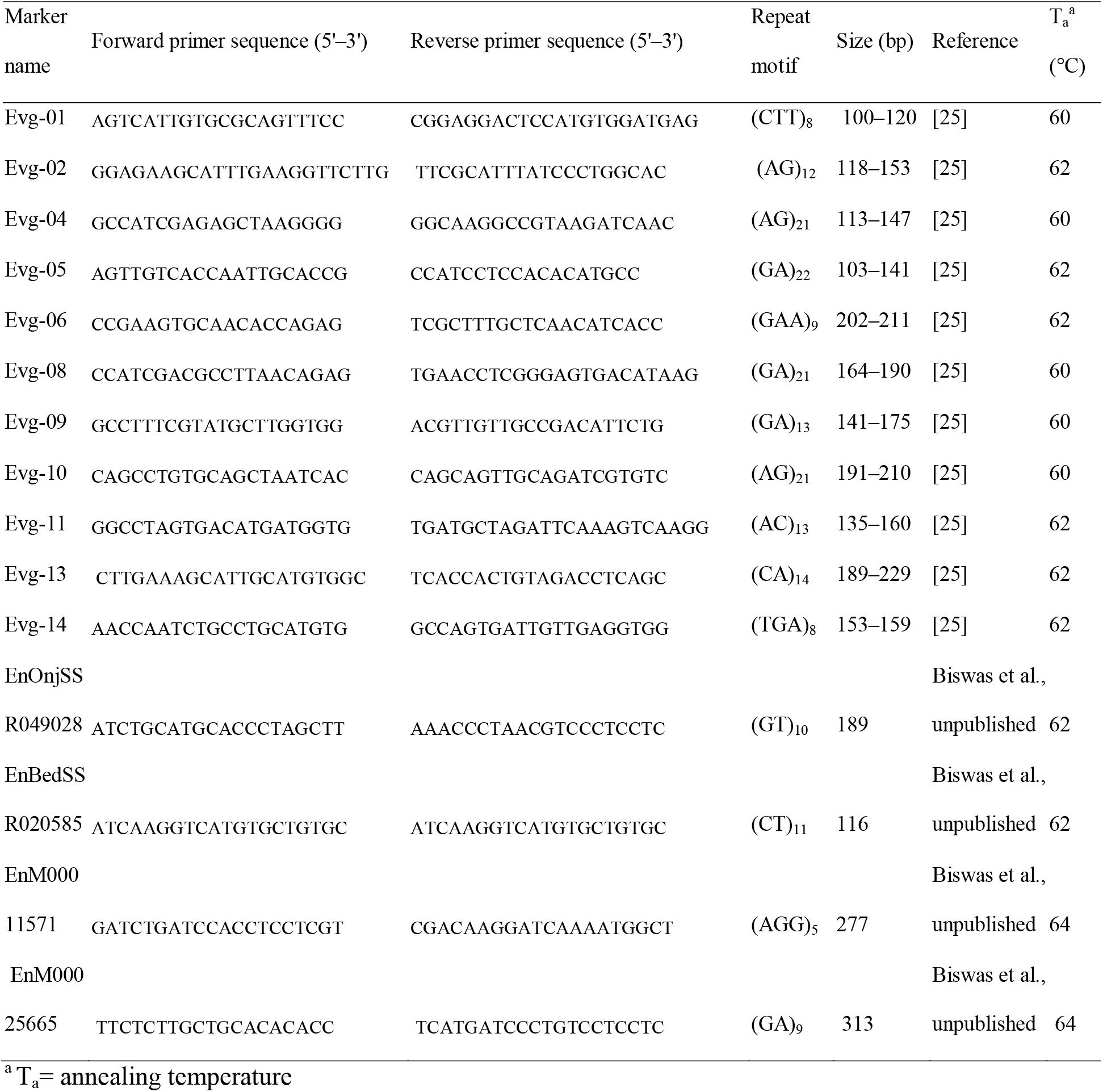
Description and source of the 15 SSR primers used in genetic diversity of enset landraces

PCR amplification was carried out in a 20 μl reaction volume containing 1.5 μl (100 mM) template DNA, 11.5 μl molecular reagent water, W 4502 (Sigma, St. Louis USA) 0.75 μl dNTPs (10 mM), (Bio line, London) 2.5 μl Taq buffer (10 × Thermopol reaction buffer), 1.25 μl MgCl_2_ (50 mM), 1 μl forward and reverse primers (10 mM) and 0.5 μl (5 U/μl) BioTaq DNA polymerase (Bioline, London) and amplified using PCR thermal cycler (BiometraTOne, Terra Universal, Germany). A three step amplification program consisted of: initial (1) denaturation for 2 min at 95 °C, (2) 35 cycles of denaturation at 95 °C for 1min, annealing at a temperature specific to each primer set (Table 2), for 1 min, extension at 72°C for 1min and (3) final extension at 72°C for 10 min. PCR products were stored at 4°C until electrophoresis.

Separation of the amplified product was accomplished in a 4% (w/v) agarose (Bioline, London) gel in 1% (w/v) TAE (Tris-acetate-EDTA) buffer containing ethidium bromide, and electrophoresed at 80 V for 3 hours. A standard DNA ladder of 100 bp (Q step 2, Yorkshire Bioscience Ltd, UK) was loaded together with the samples to estimate molecular weight. The band pattern was visualized using gel documentation system (NuGenius, SYNGENE, Cambridge, UK), and the pictures were documented for scoring.

The sizes of clearly amplified fragments were estimated across all sampled landraces. Number of different alleles (N_a_), the effective number of alleles (N_e_), Shannon’s information index (I), observed heterozygosity (H_o_), expected heterozygosity (H_e_), un-biased expected heterozygosity (uH_e_) and Fixation index for each locus were computed using GENALEX version 6.503 [27]. The Polymorphism Information Content (PIC) for each locus was computed using PowerMarker version 3.25 [28]. The N_a_, N_e_, I, H_o_, H_e_ (gene diversity), uH_e_ and Genetic differentiation (F_ST_) between the two groups of landraces were estimated using GENALEX. Analysis of molecular variation (AMOVA) was performed to evaluate the relative level of genetic variations among groups, and among individuals within a group were computed using GENALEX. The neighbor-joining (NJ) tree was constructed using the software DARwin [29] based on Nei’s genetic distance [30] to reveal the genetic relationships among the groups and individual landraces. The resulting trees were displayed using Fig Tree var.1.4.3 [31]. Principal coordinates analysis was also carried out using GENALEX, to further evaluate the genetic similarity between the landraces.

## 3. Results

Fifteen SSR markers that produced clear and scorable bands were analyzed to evaluate genetic diversity and relationship of *Ensete ventricosum* landraces used in traditional medicine and those having other use values.

### Genetic diversity

The polymorphic nature of some of the SSR markers was as shown in **Figure 2**. A total of 38 alleles were detected across 15 SSR loci in 92 genotypes (Table 4). The number of alleles generated per locus ranged from 2 to 3, with an average of 2.53 alleles. The PIC values for the markers varied from 0.16 (primer EnBedSSR020585) to 0.52 (primer Evg2) with an average of 0.41. The observed heterozygosity (H_o_) and expected heterozygosity (H_e_) ranged from 0 to 0.64 and 0.18 to 0.63 respectively, and Shannon’s information index (I) ranged from 0.31 to 1.04. The F_ST_ was significant for eight of the 15 loci (**Table 3)**.

**Figure 2.**
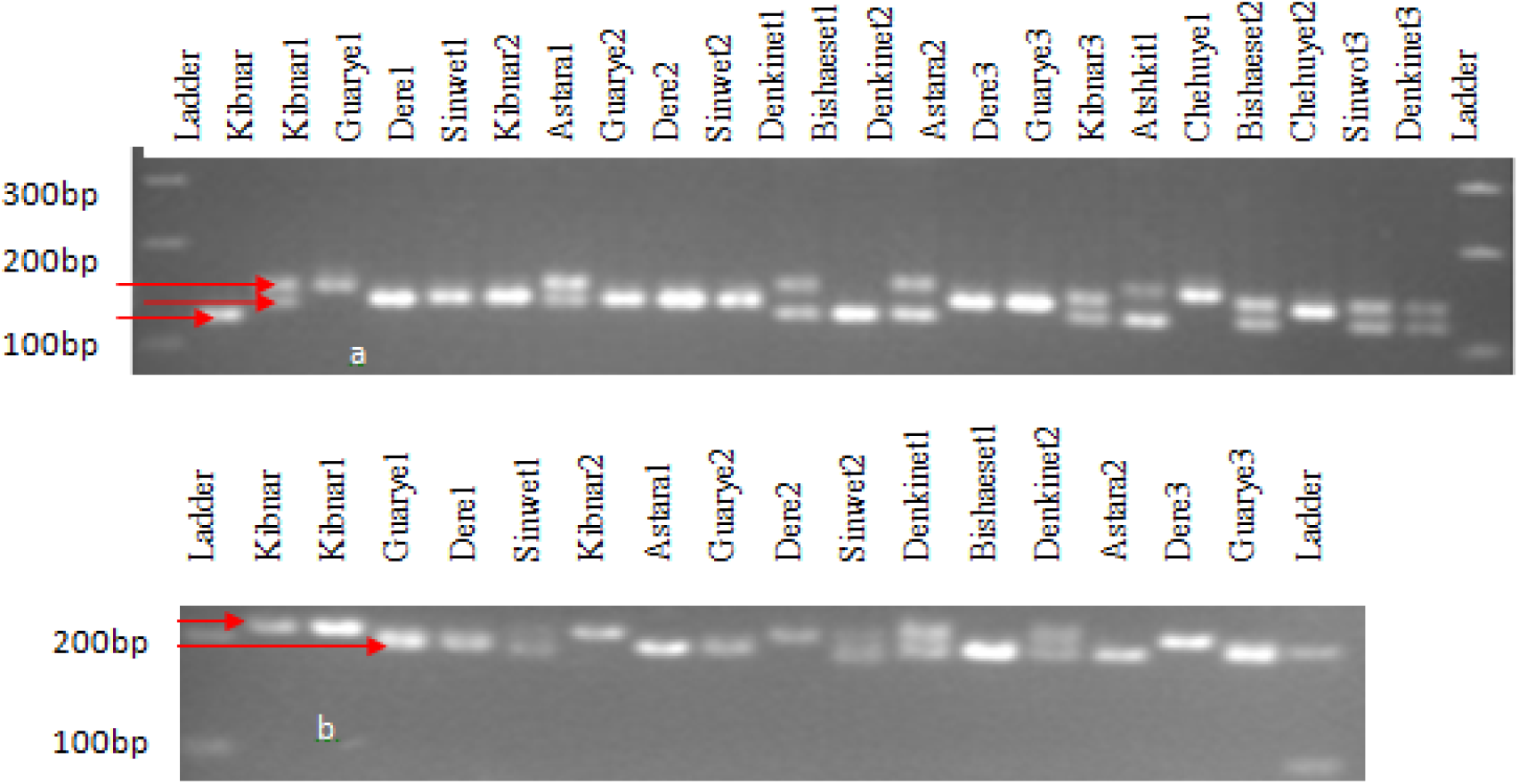
DNA fragments amplified in selected enset landraces by SSR primer; a. Evg2 (22 samples) and b. EnM00011571 (16 samples) resolved in agarose gel electrophoresis

**Table 3.** Levels of diversity indices of the SSR loci

| SSR<br>Loci | N<sub>a</sub> | N<sub>e</sub> | I | H<sub>o</sub> | H<sub>e</sub> | uH<sub>e</sub> | PIC | P<sub>HWE</sub><sup>a</sup> | F |
| --- | --- | --- | --- | --- | --- | --- | --- | --- | --- |
| Evg1 | 3.00 | 2.27 | 0.89 | 0.39 | 0.54 | 0.56 | 0.48 | 0.588<sup>ns</sup> | 0.30 |
| Evg2 | 3.00 | 2.43 | 0.97 | 0.42 | 0.59 | 0.60 | 0.52 | 0.093<sup>ns</sup> | 0.29 |
| Evg4 | 3.00 | 2.29 | 0.92 | 0.63 | 0.56 | 0.57 | 0.49 | 0.652<sup>ns</sup> | - |
| Evg5 | 2.00 | 1.82 | 0.64 | 0.54 | 0.45 | 0.46 | 0.36 | 0.006<sup>**</sup> | - |
| Evg6 | 2.00 | 1.41 | 0.43 | 0.00 | 0.27 | 0.28 | 0.26 | 0.000<sup>***</sup> | 1.00 |
| Evg8 | 3.00 | 2.21 | 0.89 | 0.51 | 0.54 | 0.56 | 0.50 | 0.000<sup>***</sup> | 0.07 |
| Evg9 | 3.00 | 2.27 | 0.93 | 0.44 | 0.55 | 0.56 | 0.49 | 0.001<sup>**</sup> | 0.20 |
| Evg10 | 2.00 | 1.82 | 0.63 | 0.00 | 0.44 | 0.45 | 0.36 | 0.000<sup>***</sup> | 1.00 |
| Evg11 | 3.00 | 1.87 | 0.74 | 0.41 | 0.46 | 0.47 | 0.43 | 0.640<sup>ns</sup> | 0.08 |
| Evg13 | 3.00 | 2.16 | 0.85 | 0.56 | 0.54 | 0.55 | 0.44 | 0.259<sup>ns</sup> | - |
| Evg14 | 2.00 | 1.92 | 0.67 | 0.64 | 0.48 | 0.49 | 0.37 | 0.017<sup>*</sup> | - |
| EnO28 | 3.00 | 2.70 | 1.04 | 0.58 | 0.63 | 0.64 | 0.59 | 0.000<sup>***</sup> | 0.08 |
| EnB85 | 2.00 | 1.23 | 0.31 | 0.21 | 0.18 | 0.18 | 0.16 | 0.814<sup>ns</sup> | - |
| EnM71 | 2.00 | 1.91 | 0.67 | 0.51 | 0.48 | 0.49 | 0.36 | 0.006<sup>**</sup> | - |
| EnM65 | 2.00 | 1.60 | 0.56 | 0.41 | 0.38 | 0.39 | 0.31 | 0.987<sup>ns</sup> | - |
| Mean | 2.53 | 1.99 | 0.74 | 0.42 | 0.47 | 0.48 | 0.41 | 0.271<sup>ns</sup> | 0.14 |

Na number of different alleles, Ne number of effective alleles, I Shannon's information index, H<sub>o</sub> observed heterozygosity, H<sub>e</sub> expected heterozygosity, uH<sub>e</sub> unbiased expected heterozygosity, PIC polymorphic information content, P<sub>HWE</sub><sup>a</sup> P-value for deviation from Hardy Weinberg equilibrium, EnO28EnOnjSSR049028, EnB85 EnBedSSR020585, EnM71 EnM00011571, EnM65 EnM00025665, ns not significant, \*\*\* =P< 0.0001, \*\* = P< 0.001 and \* =P< 0.01, F fixation Index.

**Table 4.**
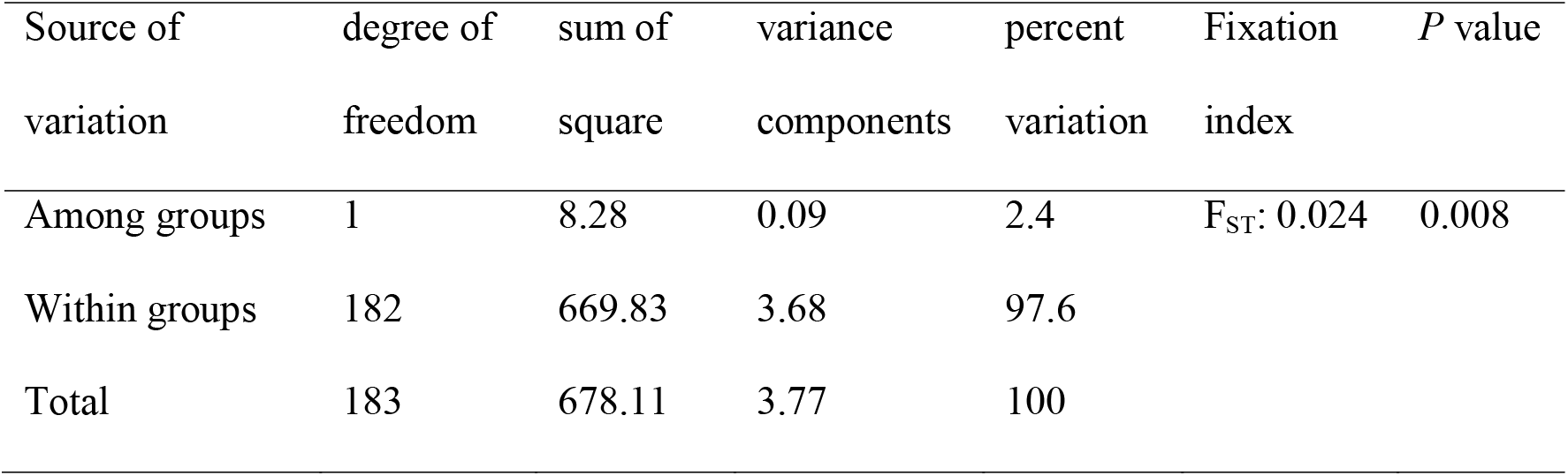
Analysis of molecular variance and fixation index for landraces used in traditional medicine and those having other use values based on data from 15 loci

| Source of variation | degree of freedom | sum of square | variance components | percent variation | Fixation index | *P* value |
| --- | --- | --- | --- | --- | --- | --- |
| Among groups | 1 | 8.28 | 0.09 | 2.4 | $F_{ST}$: 0.024 | 0.008 |
| Within groups | 182 | 669.83 | 3.68 | 97.6 |  |  |
| Total | 183 | 678.11 | 3.77 | 100 |  |  |

### Genetic differentiation and relationships

The overall genetic divergences among the two groups of enset landraces (‘medicinal’ and ‘non-medicinal’), measured in coefficients of genetic differentiation (F_ST_) was 0.024 **(Table 4)**. The analysis of molecular variance (AMOVA) also showed that 97.6% of the total variation was assigned to among individuals with in a group; while only 2.4% variation was contributed by variation among the groups **(Table 4)**.

The unweighted neighbor-joining tree cluster analysis performed using Nei’s genetic distance grouped the landraces into 4 main clusters, Cluster A–D. Generally landraces used in the treatment of a specific disease traditionally were not grouped in to the same cluster or sub cluster; instead they were mixed with those landraces having other use values (Figure 3). Similarly, the landraces originated from each location were scattered in to all the 4 clusters (data not presented). A principal coordinate analysis (PCoA) bi-plot of the first two axes (PCoA_1_ and PCoA_2_), that accounted for 33.3 % of the total genetic variability, showed a wide dispersion of accessions along the four quadrants (Figure 4). The landraces used in traditional medicine and non-medicinal of landraces did not cluster in the plot.

**Figure 3.**
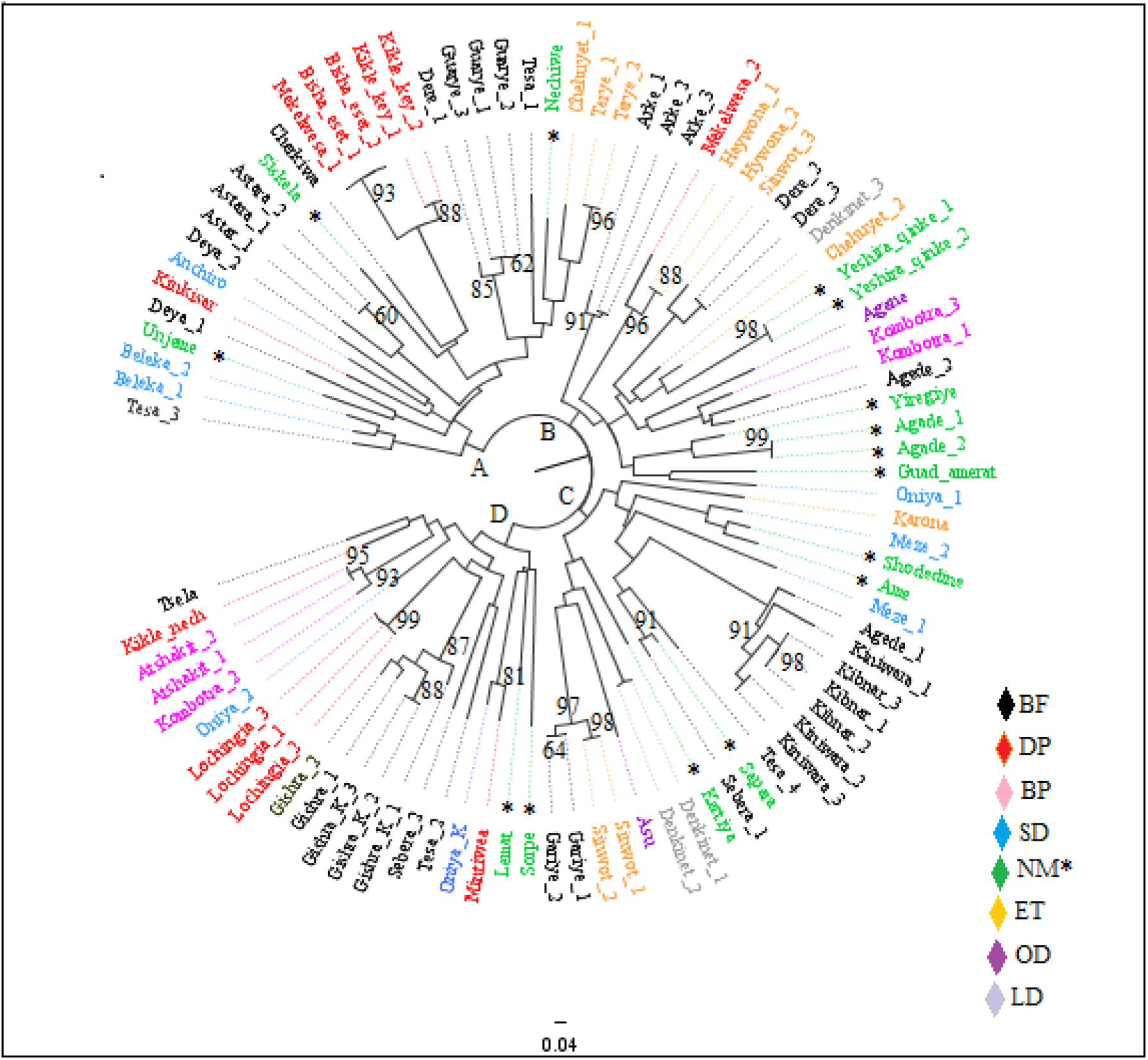
Unrooted neighbor-joining tree generated based on simple matching dissimilarity coefficients over 1000 replicates, showing the genetic relationship among 51 *Ensete ventricosum* landraces (duplicated on average 2 times) using 15 SSR markers. Landraces are color-coded according to previously identified diseases types treated by the landraces traditionally, as designated by: BF for bone fracture; DP, discharge of placenta; BP, back pain; SD, skin itching and diarrhea; NM*, ‘non-medicinal’; ET, expulsion of thorn and drainage of abscess from tissue; LD, liver disease and OD, other diseases. Numbers at the nodes are bootstrap values only > 60%

**Figure 4.**
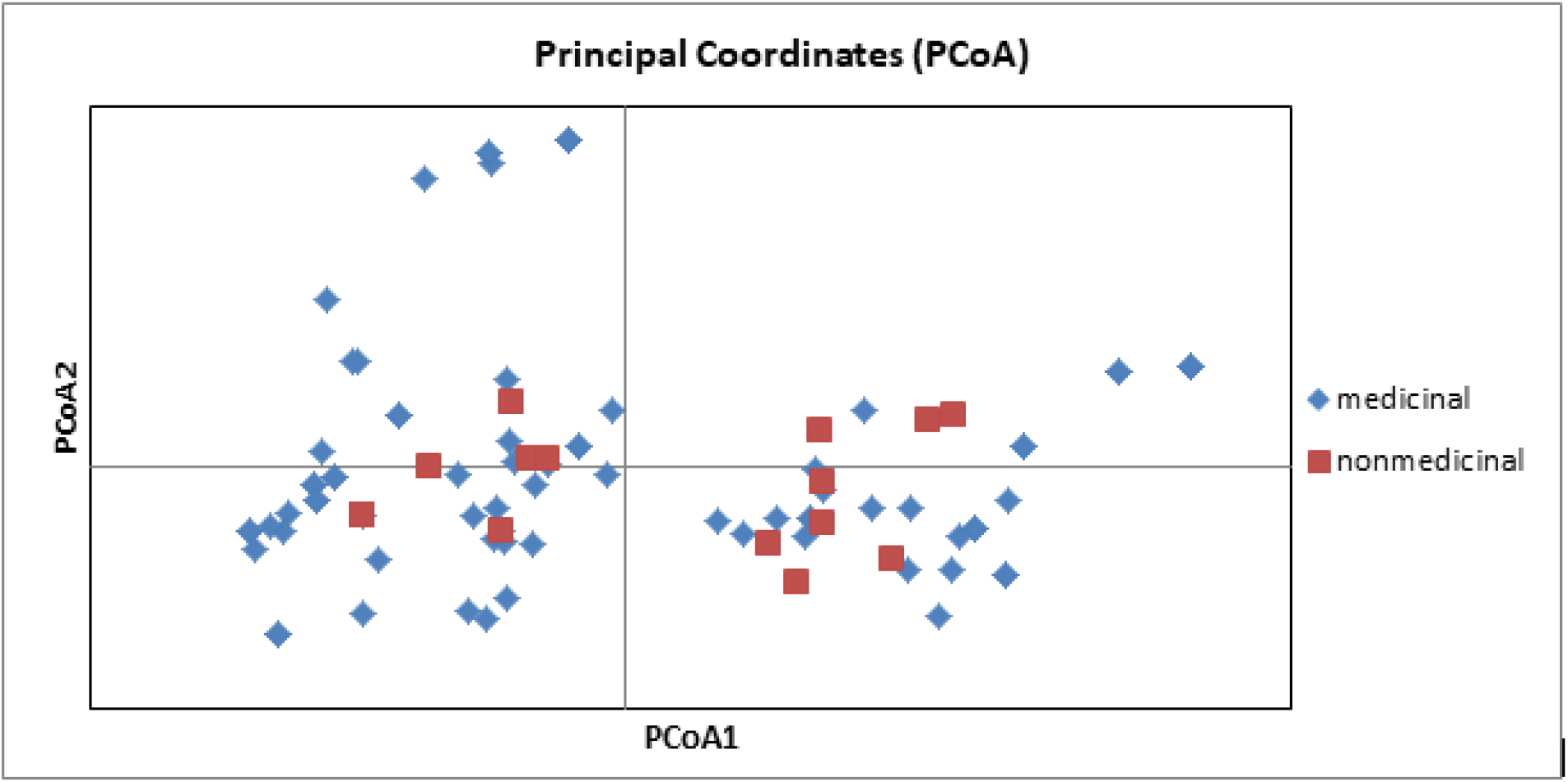
Principal coordinate analysis (PCoA) bi-plot showing the clustering of 51 enset landraces (duplicated on average 2 times) based on 15 microsatellite data, labeled based on their use value. The first and second coordinates accounting for 17.8% and 15.5% variation respectively

## Discussion

### Genetic diversity

We assessed the genetic diversity of 38 cultivated enset landraces used in traditional medicine and 13 landraces that have non-medicinal values. According to our results, moderate level of genetic diversity (H_e_ = 0.47) was detected. A relatively higher H_e_ values (0.55 and 0.59) of enset were reported [22] and [24, 25] respectively. The value of I (0.74) in the current study was also lower as compared to the earlier report (1.08) [24]. The variation probably due to the fact that our study focused on a selected group of landraces (used in traditional medicine). Lower genetic diversity estimates were reported using ISSR [21] and RAPD [20] markers. Comparisons of detailed diversity estimates from marker systems with different properties and origins of variation do not allow useful conclusions [32, 33].

### Genetic differentiation and relationships

The genetic differentiation between the landraces used in traditional medicine and those having other use values, was very low (0.024). Comparable genetic differentiation values (0.037, 0.025) among locations were reported on enset [24] and cassava [34] respectively. AMOVA also confirmed a limited (2.4%) genetic variation among the two groups of enset landrace, while the majority was contributed by variation among individual.

From the landraces that have similar vernacular names (replicate samples), the majority were closely similar genetically, and placed together in the neighbor-joining tree. This indicates that farmers have rich indigenous knowledge in identifying and naming enset landraces based on phenotypic traits, and the knowledge is shared across the growing region. However, some other replicates of landraces were placed in different cluster, indicating that genetically different landraces were given the same vernacular name. Perfect identification of genotypes using morphological traits is difficult, and the existence of homonyms has been reported previously [25].

Except two, all landraces with distinct vernacular names were found to be different, showing that vernacular names are good indicator of genetic distinctiveness in these specific groups of landraces. Whereas, the existence of 37 and 8 duplicates of landrace in diversity analysis of enset using 4 AFLP [13] and 12 RAPD[20] markers respectively, was reported. Gerura *et al*. [24], who studied 83 enset genotypes using 12 SSR markers also reported 10 duplicates of landraces. Although full identity among the landraces can only be determined when the entire genomes are compared, it is expected that the SSR markers used in the current study could sufficiently discriminate the landraces than the studies that reported higher number of duplicates. The variation of the results therefore, could be due to the sample collection method followed in the current study, which involved focusing mainly on specific landraces used in traditional medicine.

The use of some of enset landraces in traditional treatment of various human ailments in the major enset growing region of Ethiopia, SNNPR, was reported by a number of authors [8, 9, 35, 36]. However, landraces that are used in treatment of the same types of diseases did not show distinct grouping; instead landraces used to treat different diseases were mixed each other and even with those having other use values in the neighbor-joining tree, indicating that ‘medicinal’ properties do not appear to be monophyletic. Furthermore, the PCoA also showed that the two groups of landraces neither formed a separate cluster, nor did one group show greater spread or genetic diversity. From these results, it can be argued that landraces that are used in traditional medicine are not genetically distinct from other landraces.

There are several possible explanations for these observations. First, all enset landraces may have a degree of medicinal value, but specific genotypes are preferred for phenotypic or cultural reasons. Second, medicinal value may arise through genotype-environment interactions or management practices specific to those landraces i.e. they may have non-differentiated genotypes, but in situ they generate a unique biochemistry with medicinal properties. Thirdly, a number of important medicinal landraces may have been omitted, or medicinal value incorrectly assigned to non-medicinal landraces. This could serve to hinder our analysis, and make it more difficult to detect real genetic differentiation. This would also be an indication of a decline in the quality of indigenous knowledge. We also note that it is unlikely that the strong trust of society upon these landraces could not be developed after a very long period of use, and we have observed remarkable similar enset medicinal claims across a wide variety of distinct ethnic groups in multiple languages. Moreover, anti-bacterial and anti-fungal activities of a compound extracted from the unspecified *Ensete ventricosum* landrace [37], and a report [38] on medicinal property of a related species, *E. superbum*, justifies that at least some of enset landraces have real medicinal property.

## Conclusion

The study indicated the existence of moderate level genetic diversity among enset landraces used in traditional medicine. The majority of the variation was contributed by variation among individuals, indicating low genetic differentiation among the groups. Except two, all the landraces with distinct vernacular names were found to be genetically different. The landraces were not clustered based on their use values, showing no evidence for genetic differentiation between landraces used in traditional medicine and those having other use values, and the range of diversity in medicinal landraces was little different from that of landraces cultivated for food. In the future, we suggest biochemical comparison of enset landraces would complement our analysis, while genetic mapping and genome-wide association studies (GWAS) has the potential to identify genomic regions and genes associated with medicinal traits. The information from this study will be useful for identification and conservation of enset landraces used in traditional medicine, and it can provide baseline information that promotes further investigations in exploiting the medicinal value of these specific landraces.

## Additional files

**Additional file 1:** A figure showing a selective attack of enset landraces used in traditional medicine by wild animals

**Additional file 2:** Principal coordinate analysis bi-plot of the first two axes (PCoA1 and PCoA2), showing the colored data points with their corresponding landraces

## Abbreviations

AFLP: Amplified fragment length polymorphism
AMOVA: Analysis of molecular variance
CTAB: Cetyltrimethylammonium bromide
GCRF: Global Challenges Research Fund
ISSR: inter simple sequence repeat
PCoA: Principal coordinates analysis
RAPD: Randomly amplified polymorphic DNA
SNNP: Southern Nations, Nationalities, and Peoples’
SSRs: Simple sequence repeats

## Acknowledgement

This work was supported by Addis Ababa University [PhD studentship to Gizachew Woldesenbet Nuraga] and the GCRF Foundation Awards for Global Agricultural and Food Systems Research, entitled, ‘Modeling and genomics resources to enhance exploitation of the sustainable and diverse Ethiopian starch crop enset and support livelihoods’ [Grant No. 7 BB/P02307X/1]. The authors thank Ms. Hawi Niguse and Mr. Shiferaw Alemu for their assistance during DNA extraction. Dr. Fikadu Gadissa and Mr. Umer Abdi are appreciated for their help in the data analysis.

## Availability of data and materials

A figure showing selective attack of ‘medicinal’ enset landraces by rodents (porcupine and wild pig) is provided in additional file 1. Principal coordinate analysis bi-plot of the first two axes (PCoA1 and PCoA2), showing data points with their corresponding landraces is provided in Additional file 2.

## Authors’ contributions

GW, TF, KT and SD designed the experiment. GW carried out the sample collection, laboratory work and manuscript writing. GW and MK conducted data mining and carried out the data analysis. GW, JB, TF and PH are major contributors in interpreting the data. All co-authors participated in revising the manuscript and approved the final manuscript

## Ethics approval and consent to participate

Not applicable

## Consent for publication

Not applicable

